# Neuronal sex identity shapes social buffering of defensive freezing behaviour in fruit flies

**DOI:** 10.64898/2026.10.07.756549

**Authors:** Mirjam Heinemans, Rui M Gonçalves, Marta A Moita, Clara H Ferreira

**Affiliations:** Champalimaud Neuroscience Programme, Champalimaud Research, Champalimaud Foundation, Avenida de Brasilia, Lisbon, Portugal; School of Geography and Natural Sciences, Ellison Building, Northumbria University, Newcastle upon Tyne, NE1 8ST, UK

## Abstract

Sex differences are pervasive across the animal kingdom^1,2^, shaped by reproductive strategies, ecological roles, and physiological demands. Neurobiological studies have largely focused on sex differences in reproduction and aggression^3,4^, leaving understudied how sex shapes other survival-relevant behaviours such as responses to predatory threats. Previous work in vertebrates uncovered sex differences in individual defence responses deployed by rats^5^ and in social modulation of freezing responses in zebrafish^6^. The origin of these differences and their potential underlying neuronal circuitry remain however unknown. Here we characterise defensive responses to a visual threat of female and male fruit flies tested alone and in groups. Individually, defensive responses are comparable across sex, with similar freezing levels and moderately increased escape responses in males. In groups, however, we uncovered strong sex differences in freezing as males show consistently shorter freezing bouts, reflecting a stronger social buffering effect than in females. Using a generalised linear mixed model, we demonstrate that the shorter freezes of male flies, when tested in groups, are not explained by well-known locomotor differences between sexes^7,8^. Although males move more and faster, generating stronger social motion cues that mediate the ‘safety in numbers’ effect in female flies^9,10^, their response to the surrounding motion is not increased. We further identify the nervous system as the source of this sex-dimorphic behaviour by feminising males through pan-neuronal ectopic expression of transformer, a key regulator of the cell-autonomous sex-differentiation pathway. Finally, we ruled out the visual projection neurons, lobula columnar neurons 11 (LC11), detectors of social motion^10^, as the source of the neuronal-sex differences observed. This study identifies a neuronal origin of sex differences in socially modulated behaviour that extends beyond sex-specific behavioural modules or locomotor function, important to understand differential impact on survival upon threat across sexes.

## Results and discussion

### Defensive responses in individually tested flies are overall similar across sex

To assess potential sex differences in defence behaviours in response to visual looming stimuli (Figure 1A) we first analysed the individual behaviour repertoire of females and males. We found overall similar behavioural patterns across sex with predominant freezing responses during the stimulation period, interspersed with evasive manoeuvres, namely jumps and more rarely escape runs characterised by speed increases in response to looms (Figure 1B). Both females and males predominantly initiated freezing responses within the loom presentation window (Supplementary Figure 1A) and show a marginal difference in the proportion of time spent freezing during the stimulation period (Figure 1C, PropF_F_ = 0.92, IQR 0.80–0.96, PropF_M_ = 0.87, IQR 0.69–0.94, δ = -0.17, 95% Bias-Corrected and Accelerated bootstrap confidence intervals, CI [-0.33, -0.01], p = 0.03). A closer inspection of freezing behaviour failed to show significant differences in the length of freezing bouts across sex, though females showed a longer tail (Figure 1D, median Fbout_F_ = 5.50 s, IQR 1.34-44.75, Fbout_M_ = 7.13 s, IQR 2.16-24.78, Anderson-Darling test, AD = 1.06, p=0.12; n_F_ = 255 bouts, n_M_ = 304 bouts). As with freezing responses, the timing of jump initiation upon a loom was similar across sexes (Supplementary Figure 1B), but males jumped more throughout the 5 minutes of stimulus presentations (Figure 1E, NJumps_F_ = 1, IQR 0–1, NJumps_M_ = 2, IQR 0.50–4, δ = 0.41, 95% CI [0.26, 0.55], p < 0.001). Finally, while individually tested males showed higher walking speeds during the baseline (Figure 1F, SpeedBase_F_ = 11.56 mm/s, IQR 10.71–12.75, SpeedBase_M_ = 13.65 mm/s, IQR 12.70–14.59, δ = 0.66, 95% CI [0.53, 0.77], p < 0.001) and stronger speed increases relative to speed prior to the loom (Figure 1G, SpeedInc_F_ = 2.65 mm/s, IQR 0.94–3.72, SpeedInc_M_ = 3.92 mm/s, IQR 2.07–8.24, δ = 0.36, 95% CI [0.09, 0.58], p = 0.006; Supplementary Figure 1C), they did not show an increased probability of speeding up upon a looming stimulus, which was low for both sexes (Supplementary Figure 1D, ProbSpeedInc_F_ = 0, IQR 0–0.05, ProbSpeedInc_M_ = 0, IQR 0–0.05, δ = 0.10, 95% CI [−0.04, 0.25], p = 0.13). In summary, in our experimental conditions, where freezing behaviour dominates, individual defensive responses to looming stimuli were grossly similar across sexes. While differences in freezing were marginal, escape responses were increased in males, both through increased number of jumps and more vigorous walking speed increases.

**Figure 1.**
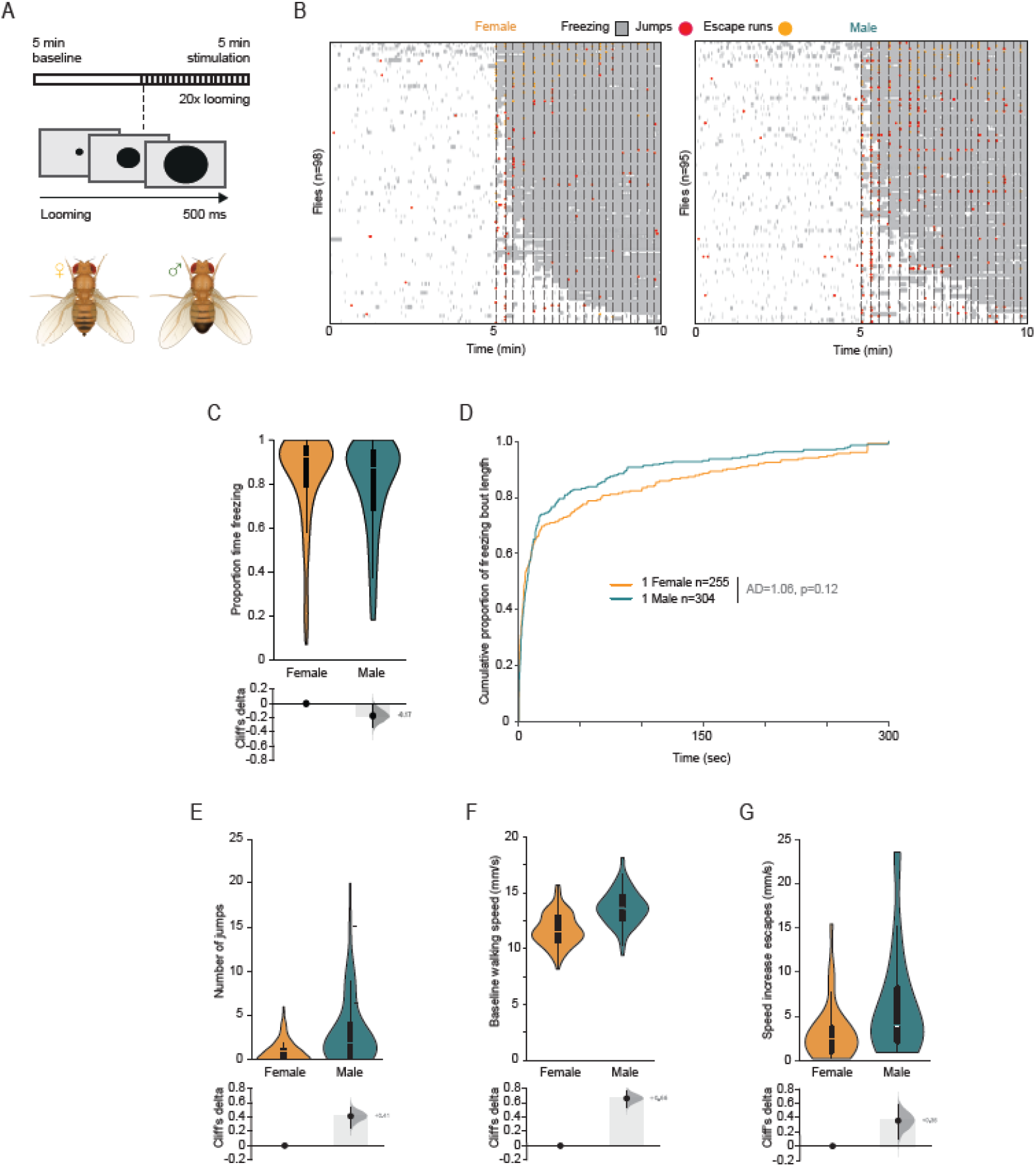
Individual defensive responses per sex. **(A)** Protocol: we imaged single sex flies in arenas exposed to a 5-min white screen followed by twenty 500 ms looming presentations, every 10–20 s. **(B)** Defensive responses throughout the experiment: freezing (grey), jumps (red) and escape runs (purple) for control females (left, n=98) and male flies (right, n=95). Each row corresponds to one fly, rank-ordered by maximum time freezing during stimulation. **(C)** Proportion of time freezing throughout the stimulation period. (**D)** Cumulative distribution of freezing bout length for females (orange) and males (teal). Anderson-Darling (AD) test statistic and p-value are indicated. **(E)** Number of jumps in response to looming stimuli. **(F)** Average walking baseline speed. **(G)** Speed increase during escape runs in response to looming stimuli. **(C, E-G)** Top: Violin plots representing the probability density distribution of individual fly data bound to the range of possible values, with boxplot elements (central white dot, median; box limits, upper (75) and lower (25) quartiles; whiskers, 1.5 × interquartile range). Bottom: Cliff’s delta effect size for the female–male comparison (black dot, observed Cliff’s delta; grey curve, bootstrap sampling distribution; vertical line, 95% confidence interval).

### Groups of males show stronger social modulation of freezing bout length

Since the social environment is a strong modulator of defensive responses, often leading to social buffering, that is decreased responses to threat in line with the ‘safety in numbers ‘effect^11–13^, we next analysed loom-triggered behaviours in groups of flies. We previously found that female flies display social ‘buffering’ of freezing decreasing this response with increasing levels of motion by neighbouring flies^9,10^. In this study we tested either male or female flies in groups of up to five, comparing freezing behaviour across sex for each group size. First, we found that males also showed an incremental decrease in freezing responses with increasing group sizes, establishing social buffering of freezing in male flies (Figure 2A, Supplementary Tables 1a and 1b). The decrease in freezing appeared to be stronger than that observed in females when we compared the proportion of time freezing across sex (Supplementary Figure 2A, Supplementary Table 2). Indeed, although the probability of entering freezing remained largely the same regardless of social context (Supplementary Figure 2B, Supplementary Table 3), unlike in individually tested flies (Figure 1D), males in groups systematically exhibited shorter freezing bouts when compared to those of same-sized groups of females, suggesting a stronger social modulation in males (Figure 2B, Supplementary Table 4).

**Figure 2.**
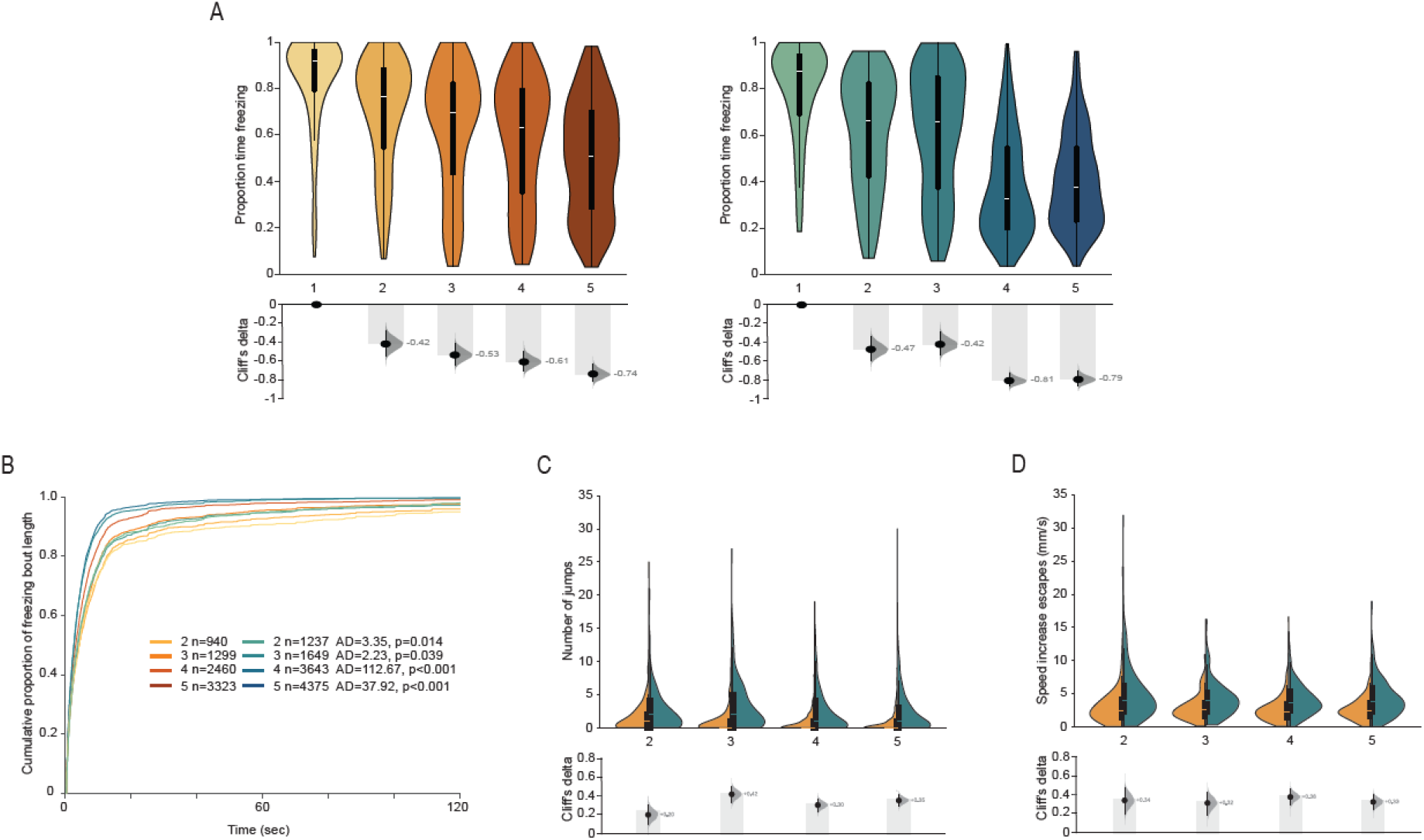
Group defensive responses per sex. **(A)** Proportion of time flies spend freezing for females (left) and males (right) tested individually and in groups of two to five flies. **(B)** Cumulative proportion of freezing bout length for females (shades of orange) and males (shades of teal), separated by group size (2–5, see legend inset for colour code and number of freezing bouts). Anderson-Darling (AD) test statistic and p-value are indicated for each group size. **(C, D)** Number of jumps **(C)** and speed increase during escape runs **(D)** in response to looming stimuli, shown as split-violin plots per group size (2–5; females, orange; males, teal). **(A, C, D)** Top: Violin plots representing the probability density distribution of individual fly data bound to the range of possible values, with boxplot elements (central white dot, median; box limits, upper (75) and lower (25) quartiles; whiskers, 1.5 × interquartile range). Bottom: Cliff’s delta effect size for individual-group or the female–male comparison (black dot, observed Cliff’s delta; grey curve, bootstrap sampling distribution; vertical line, 95% confidence interval).

Sex differences in escape responses were similar to those observed in individually tested flies (Figure 1E and 1G) and maintained across group size (Figure 2C, Supplementary Table 5), with males showing more jumps and stronger speed increases than females (Figure 2D, Supplementary Table 6) with no consistent change in escape run probability (Supplementary Table 7). In conclusion, sex differences were more prominent in threatened groups of flies, as differences in freezing become apparent only when tested in groups, with males freezing less than females, while sex differences in escape responses were maintained. Importantly, the sex differences observed do not seem to reflect a trade-off between flee or freeze strategies, since the decrease in freezing in groups of males resulted from a decrease in freeze bout duration with negligible effects on probability of responding to the loom with freezing, and escape differences were similar across social contexts.

### Sex differences in freezing bout length are independent of sex differences in locomotion and have a neuronal origin

Having identified sex differences in loom-triggered freezing, which remains the dominant defensive response of flies in groups despite the social buffering effect (individuals PFentry_F_ = 1.00, IQR 0.43-1.00, PFentry_M_ = 1.00, IQR 0.50-1.00; groups of 2 to 5 PFentry_F_ = 0.67-0.75, IQR 0.43-1.00, PFentry_M_ = 0.67-0.75, IQR 0.47-1.00; Figure 2, Supplementary Figure 2), we next set out to uncover the source of these differences, resorting to generalised linear mixed models. Given the well-documented modulation of predator-prey interactions by locomotor cues, such as the speed of the predator and prey, as well as movement of surrounding animals (reviewed in^14,15^; in flies^10,16^), and the established locomotor sexual dimorphism in fruit flies (see references^7,17^), we hypothesised that features of locomotion could explain the observed sex differences in freezing. Social buffering affected mostly the duration of freezing responses (Figure 2B), therefore we modelled the distribution of freezing bout durations. As predictors we used variables characterising the locomotion of flies, such as the speed of flies before each loom (pre-loom speed, previously identified as a modulator of individual freezing behaviour^16^), baseline walking speed and proportion of time walking (locomotor patterns previously shown to differ across sex^7,17^), social motion (a measure of the amount of motion a fly is exposed to while freezing, caused by the movement of surrounding flies, that leads to decreased freezing^9,10^, see methods), and finally loom number (see methods). We used a log-normal distribution as this best fit our data (Supplementary Table 8) with fly identity and group identity as random effects. For a female fly at mean values of all continuous predictors, the model predicted a geometric mean freezing bout duration of 5.2 s. This analysis uncovered a prominent sex effect on freezing bout length (males displayed 29.1% shorter bouts than females) that surpasses any interaction of sex with relevant predictors (Figure 3A; Supplementary Table 9; Supplementary Figure 3A and B).

**Figure 3.**
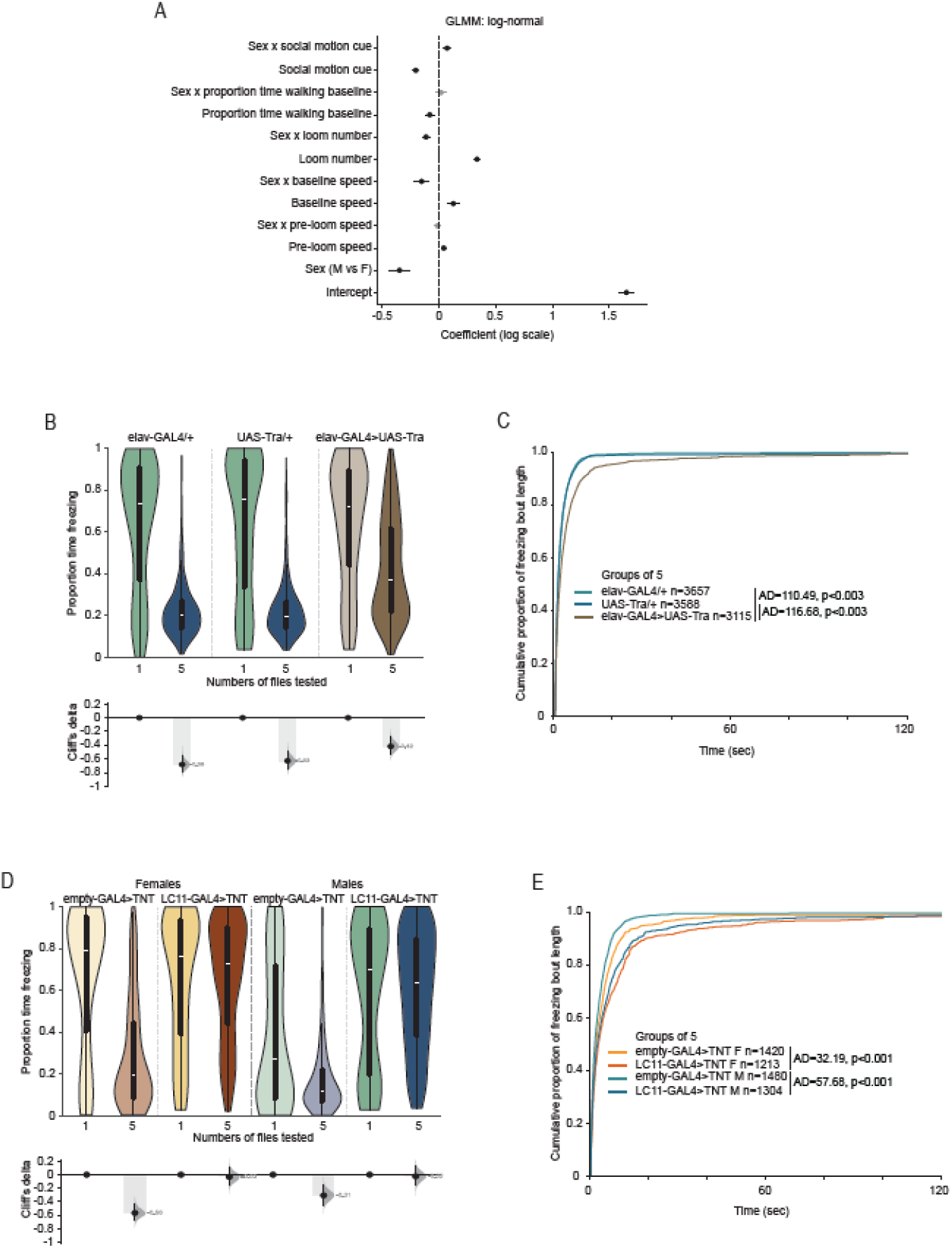
Sex differences in freezing bout length surpass individual and social predictors and are neuronally driven. **(A)** Generalised linear mixed model of freeze bout duration, with a log-normal distribution and log link, with sex as a categorical predictor (female as the reference level) and standardised continuous predictors. Posterior mean coefficients (dots) and 89% credible intervals (lines; black excludes zero; grey crosses zero). **(B-E)** Proportion of time freezing during the stimulation for individuals and groups of 5 **(B, D)** and cumulative proportion of freezing bout length in groups of 5 **(C, E)** for pan-neuronally feminised males (*elav-GAL4>UAS-Tra*) and parental controls (*elav-GAL4/+; UAS-Tra/+*) **(B, C)** and LC11 silenced neurons (*LC11-GAL4>TNT*) and control (*empty-GAL4>TNT*) **(D, E)** (see legend inset for colour code and number of flies tested). Anderson-Darling (AD) test statistic and Bonferroni-corrected p-value are indicated.

The strongest positive predictor of freeze bout length was loom number, followed by baseline speed and pre-loom speed (40.1%, 13.8% and 4.6%, respectively, Figure 3A). Conversely, social motion cues had the strongest negative impact on freezing durations, followed by the proportion of time walking during the baseline (18.1% and 7.5%, respectively, Figure 3A). This means that the more looms the flies were exposed to and the faster flies moved, the longer the freeze bouts, whereas flies exposed to stronger social motion cues while freezing and flies that walked more during the baseline showed shorter freezes. Some of these predictors showed an interaction with sex. Male flies showed attenuated positive modulation by loom number and baseline speed, as well as attenuated negative modulation by social motion cues (Figure 3A). The aforementioned model included all freeze bouts, however in a minority of cases (∼10% overall; Supplementary Table 10) flies showed very long freeze bouts that extended past the next, or multiple, looms. Interpreting the duration of such bouts may be more difficult, therefore we ran the same model including only the bouts contained within the interval between looms (the majority of cases), that is the ones where the fly resumed moving before the next loom. Again, we found a positive main effect of loom number and movement speed of the focal fly both during the baseline period and immediately prior to the loom, and a negative effect of sex and social motion around the fly, maintaining the attenuation of the impact of baseline walking speed on freeze bout duration of male flies (Supplementary Figure 3C). Both models confirm the decreased freezing of males when exposed to threat in groups, however contrary to our expectation male flies did not show increased sensitivity to social motion cues, the main mediator of the social buffering effect on freezing. It could still be possible that the shorter freeze bouts observed in males resulted from the increased social motion cues male flies were exposed to while in groups, since male flies walk more and faster than females. To test this possibility, we simulated freeze bout durations of male flies (using the predictor coefficients estimated for males, that is the weight of each predictor and their interactions) but using the values of female locomotion (baseline and pre-loom speeds as well as social motion cues), as a control we simulated male freeze bout duration with their own locomotion values (Supplementary Figure 3D). The same simulations were run for female flies. We found that simulated males yielded similar freeze bout durations regardless of the locomotion features used in the model (whether male or female) and that both were shorter than the simulated female freeze bout durations (Supplementary Figure 3D; Supplementary Table 11). In the case of females, a significant difference was found when using male or female locomotion values, reflecting even further increased freezing bouts with the male values, possibly because of the female higher sensitivity to baseline speed (Supplementary Figure 3D). This finding indicates that the sex difference in freezing bout length reflects differences in how each sex responds to the predictor variables, rather than differences in the predictor values themselves. Finally, analysing groups of three flies with different proportions of same sex flies confirmed the sex difference persists in mixed groups, indicating it is an inherent characteristic of the sex of the focal animal analysed (Supplementary Figure 4A, B and C, Supplementary Tables 12-14). In conclusion, this modelling analysis supports a general role for sex in dictating freezing bout differences that are largely independent of individual and social locomotion-related predictors. These findings are unexpected since the increase in locomotion of male flies, resulting in stronger social motion cues in groups of male flies, does not explain the stronger social buffering of freezing in males.

Given that the simple explanation of stronger male social buffering of freezing due to higher locomotion was not verified, we next explored the neuronal origin of this effect, as previous work identified sex differences in neuronal activity levels, modulation or circuitry underlying reproductive and aggressive behaviours, locomotor patterns and exercise physiology^7,8,18–20^. To explore this possibility, we exploited the cell-specific sex differentiation pathway by manipulating *transformer*, a key regulator of the cell-autonomous sex-differentiation pathway^21,22^. We drove ectopic expression of *transformer* using a pan-neuronal driver to feminise males. Freezing in groups was strongly affected by switching the sex of the nervous system, with groups of feminised males (*elav-GAL4>UAS-Tra*) showing lower social buffering of time spent freezing (Figure 3B, Supplementary Table 15) and longer freezing bout length in groups compared to parental controls (Figure 3C, Supplementary Table 16). Since we had previously identified LC11 neurons^10^, responsive to small moving objects^23,24^, as mediators of the visual social motion cues in females, we tested whether the sex differences in social buffering of freezing resulted from differential requirement of these neurons across sexes. To this end, we silenced LC11 neurons in male flies, using a *LC11-GAL4* line to express tetanus toxin light chain (TNT), which cleaves neuronal synaptobrevin preventing synaptic release of neurotransmitter^25^. In line with the GLMM analysis, we found that the sex difference in the magnitude of the social modulation of freezing effect is not due to the visual perception of social motion cues, as constitutively silencing LC11 neurons (*LC11-GAL4>TNT*) leads to lowered social buffering of time spent freezing (Figure 3D, Supplementary Table 17) and longer freezing bout length in groups (Figure 3E; Supplementary Table 18) compared to controls (*empty-GAL4>TNT*) in both sexes, while sex differences prevail across genotypes (Supplementary Table 19). In conclusion, the sex differences observed in freezing bout length in groups are driven by the sexual identity of the nervous system and do not reflect processing differences at the level of visual projection neurons.

## Conclusions

Our work uncovered sex differences in the deployment of defensive responses to a threatening looming stimulus mimicking a predator attack. Unlike previous work in rats^5^, where females displayed more darting behaviour, while males showed more freezing in an auditory cued fear conditioning task, we found that males and female flies deploy the same type of defensive strategies to looming stimuli, which are similar in magnitude for freezing yet higher in males for number of jumps and in the magnitude of speed increase in escape runs. Zebrafish, on the other hand, have been found to display sex difference in socially modulated freezing responses^6^, with males showing higher levels of ‘social buffering’, a decrease of freezing responses in groups, but also higher contagion, increased freezing responses to freezing of conspecifics exposed to an olfactory alarm signal. This heightened social attunement in males is thought to reflect broader life-history strategies involving higher risk taking and energetically demanding inter-sex competition which increase predator encounters and allow rapid control of fear responses to save energy in absence of a predator and increase responsiveness when it is present. Similarly, we find that male flies display lower freezing levels in groups, which due to its metabolic cost^26^, could also potentially reflect similar life-history trade-offs in resource competition.

We also identified that the male freezing modulation by the social environment is an inherent property of being male that does not arise from the variables we have experimental access to in our behavioural paradigm. Previous work had already identified sex differences in locomotion^7,8,17^ and we identified that males walk at higher speeds than females, and also show higher number of jumps and higher magnitude escape runs. These differences in individual locomotion which impact freezing responses^16^ and also the level social motion cues which have a direct effect on freezing responses^9,10^ do not however explain the sex differences in freezing bout length as we found through our GLMM analysis. The neuronal basis of the sex differences we observe in freezing bout is hence unlikely to be the same source as that of locomotor sex differences^7,8^.

We further established that the sex differences in freezing bout length are neuronally controlled as feminising the male nervous system increased freezing durations, yet they are unlikely to be instantiated at the level of LC11 neurons we previously found to mediate the regulation of freezing by social motion cues in females^10^. One possibility would be that males and females rely on different visual projection neurons to detect visual motion, however, inactivating these neurons in both females and males decreases the social buffering effect on freezing. This raised the possibility that differences in sensitivity of these neurons to social motion cues could explain the sex differences found. However, our model clearly demonstrates that male flies are not more sensitive to the social motion cues. In addition, the stronger social motion cues males are exposed to do not explain the differences in freeze bout durations.

These results point to an interesting common processing of social motion cues yet an independent sex-specific modulation of group behaviour. Consistent with this, visual projection neurons, including the LC neurons, are almost entirely isomorphic between the sexes in morphology and connectivity, with dimorphism arising downstream, in central brain circuits^27^. Previous work in reproduction, aggression and social modulation of locomotion in males converges on internal-state-dependent activity of P1 neurons^8,28^, in some cases acting through modulation of visual projections LC10a^29,30^. Females have a related sex-dimorphic circuit, with pC1d/e neurons at its centre, which act upstream of aIPg, a node recently shown to shape the flow of visual information carried by LC9, LC10a, and LC11^31^. It will be interesting to pursue whether the differences in freezing in a group setting are also mediated via these neurons, which could make them a pervasive hub in sex differences in behaviour. More broadly, our work sets the scene to uncover detailed neuronal underpinnings of socially-modulated, sex-specific behaviour beyond the canonical domain of reproduction and aggression, in an equally important behaviour for survival, the response to a predatory threat.

## Authors ‘contributions

M.H.: investigation, data curation, software, visualisation. R.G.: investigation, data curation. M.A.M.: funding acquisition, supervision, methodology, writing – review and editing. C.H.F.: conceptualisation, funding acquisition, supervision, data curation, formal analysis, investigation, methodology, software, visualisation, writing – original draft, writing – review and editing.

## Acknowledgements

We thank the Fly Platform at the Champalimaud Centre for the Unknown for providing fly work related infrastructure, expertise, and support; Juan Castiñeiras de Saa for helpful discussions regarding the application of generalised linear mixed models; the Moita and Huetteroth lab members for helpful discussions of the data and feedback on the paper. Stocks obtained from the Bloomington *Drosophila* Stock Center (NIH P40OD018537) were used in this study.

## Methods

### Fly lines and husbandry

Flies were reared on standard fly medium and kept at 25°C and 70% humidity in a 12:12 h dark:light cycle. Experimental animals were mated females and mated males, tested only once when 4–7 days old.

Wild-type flies used were Canton-S. *LC11-GAL4* (*w[1118]*; *P{y[+t7*.*7] w[+mC] = GMR22H02-GAL4}attP2* RRID:BDSC_49304) and *empty-GAL4* (*w[1118]; P{y[+t7*.*7] w[+mC]=GAL4*.*1Uw}attP2* RRID:BDSC_68384) were obtained from the Bloomington *Drosophila* Stock Centre. *UAS-TNT* (*UAS-(+) TNT*), *UAS-Tra* (*w[1118]; P{w[+mC]=UAS-tra*.*F}20J7*) and *elav-GAL4* (*w[1118]; elav-GAL4*) lines were obtained from the Chiappe, Vasconcelos and Rhiner laboratories, respectively. We used FlyBase^32^ to find information on stocks.

### Behavioural apparatus and experimental protocol

Behavioural experiments were performed as described in our previous studies^9,10^ (Figure 1A). Briefly, we imaged unrestrained flies in 68 mm diameter polyethylene terephthalate arenas, with slanted walls at 11° and a central flat portion of 32 mm diameter. Visual stimuli were delivered on a screen (Asus ROG STRIX XG258Q monitor running at 240 Hz, tilted 45° over the stage). The experiments ran for 10 mins comprising a 5-min baseline during which the screen was white, followed by a 5-min stimulation period where twenty 500 ms black looming stimuli were delivered randomly every 10 to 20 s (virtual object length of 1 cm, exponentially expanding at a speed of 50 cm/s; half-length to speed ratio 20 ms, corresponding to biologically relevant threats, which mimic damselfly attacks^33^). Two arenas were imaged simultaneously on a stage, backlit with infrared-light (940 nm) delivered by a custom-built LED board. Videos were acquired through Bonsai^34^ at 60 Hz and 1,280 width × 960 height resolution using two USB3 cameras (PointGrey Flea3). We tested wild-type flies alone and in groups from two to five flies, and neuronally sex-reversed flies and controls individually and in groups of five.

### Data analysis

As reported in our previous studies^9,10^, we used idTracker^35–37^ to obtain the x, y coordinates of each individual fly throughout the video, followed by processing with a custom Bonsai script to analyse pixel change in a 4 mm × 4 mm square around the centre of mass of the fly. Data were then analysed using custom scripts in Spyder (python 3.8). First, as we previously described^9,10,16^, we classified behaviours based on set values for pixel change and speed, into walking, freezing and jumping. Walking frames were defined as those with pixel change greater than zero and speed between 3.9 and 75 mm/s; stable walking bouts required a minimum engagement of 100 ms, with breaks of up to 50 ms tolerated as noise. Jumps were detected as local peaks in speed exceeding 75 mm/s, at least 3 frames apart (detect_peaks, version 1.0.4); a window of ±4–5 frames around each detected jump was excluded from other behavioural classifications to prevent jump-related motion artefacts from being misclassified.

Freezing was classified as described in our previous study^9^, a zero-pixel change detected around the fly for at least 500 ms, with a 50 ms buffer for noise. Freezing in response to looms was determined using a time window starting 500 ms before each loom until 2.5 s after the loom onset. The proportion of time freezing during the stimulation (last five min) was calculated by adding the frames in which freezing occurred and dividing this by the total duration of that period. Freezing bout lengths were calculated by measuring the time elapsed from freezing onset to offset, or until the experiment ended (when flies, once freezing, remained immobile for the rest of the test session). Since flies can respond to looms with brief startle responses that peak near the point of maximal apparent object size^16^ within an otherwise continuous freezing bout, we treated freezing bouts with interruptions exclusively during the 500 ms loom presentation window as part of the same bout rather than as a genuine bout offset, provided the fly resumed freezing within 500 ms of loom offset; freeze bout duration was determined up to the first frame at which freezing ceased.

Speed increases in response to looms were identified by comparing bootstrapped estimates of mean speed in the 1 s window preceding loom onset and the 1 s window following loom offset. For each loom, we generated 100 bootstrap resamples (with replacement, n = 30) of speed values within each window and compared the resulting distributions of resampled means using a two-sample t-test. A loom was classified as eliciting a speed increase if post-loom speed significantly exceeded pre-loom speed (t < 0, p < 0.05), the fly was not freezing during the pre-loom window, and no jumps were detected between loom onset and 1 s after loom offset. Speed increase magnitude was calculated as the difference between post- and pre-loom mean speed; speed increase probability was calculated as the proportion of the 20 looms where behaviour met these criteria.

For the generalised linear mixed model (GLMM), we calculated focal pre-loom speed, baseline speed, proportion of time walking, as well as the social motion cues flies are exposed to during a freezing bout. Focal pre-loom speed was calculated as the mean speed of the focal fly in the 500 ms immediately preceding each loom onset. Baseline speed was calculated as the mean speed of the focal fly during walking bouts occurring within the baseline period. Proportion of time walking was calculated as the fraction of frames within the baseline period classified as walking. The social motion cues to which a focal fly is exposed were calculated as previously^9,10^, as the summed product of speed and angle on the retina (θ) of a focal fly that each of the surrounding flies produces: 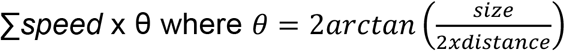 (size was held constant at 2.5 mm). Social motion cues were averaged across the duration of each freezing bout.

Statistical analysis was done resorting to the Anderson-Darling test using SciPy statistics^38^, and the Cliff’s delta effect size for non-normalized data, calculated using DaBest python package^39^. Effect sizes and 95% confidence intervals were calculated using bootstrap resampling (5000 resamples per pair). Where required p-values were adjusted via Bonferroni correction to reflect the number of comparisons.

### Generalised linear mixed model

Freeze bout duration was modelled using the Bambi^40^ python package for GLMM fitting, with fly and group identity as random intercepts, sex as a categorical predictor (female as reference level), and focal pre-loom speed, baseline speed and proportion of time walking, as well as social motion cues and loom number as continuous predictors, with sex interacting with each continuous predictor. Continuous predictors were standardised (z-scored) prior to model fitting; posterior estimates are therefore reported as the change in log(freeze bout duration) per one standard deviation change in each predictor. To select an appropriate error distribution, we compared the fit of Gamma and Weibull distributions with a log-link and a log-normal distribution using leave-one-out cross-validation (LOO-CV). The log-normal was best supported (Supplementary Table 5) and used for all models. Posterior estimates were obtained via Markov Chain Monte Carlo sampling (4 chains, 1,000 draws each following 1,000 tuning draws). Posterior estimates for all fixed effects were robust to the choice of prior, remaining qualitatively unchanged when default package priors were replaced with tighter or wider alternatives. For modelling the data of the mixed groups of 3 we used the predictors found relevant in the single sex groups and added the continuous predictor number of surrounding males.

