## Supplementary figures for "Neuronal sex identity shapes social buffering of defensive freezing behaviour in fruit flies"

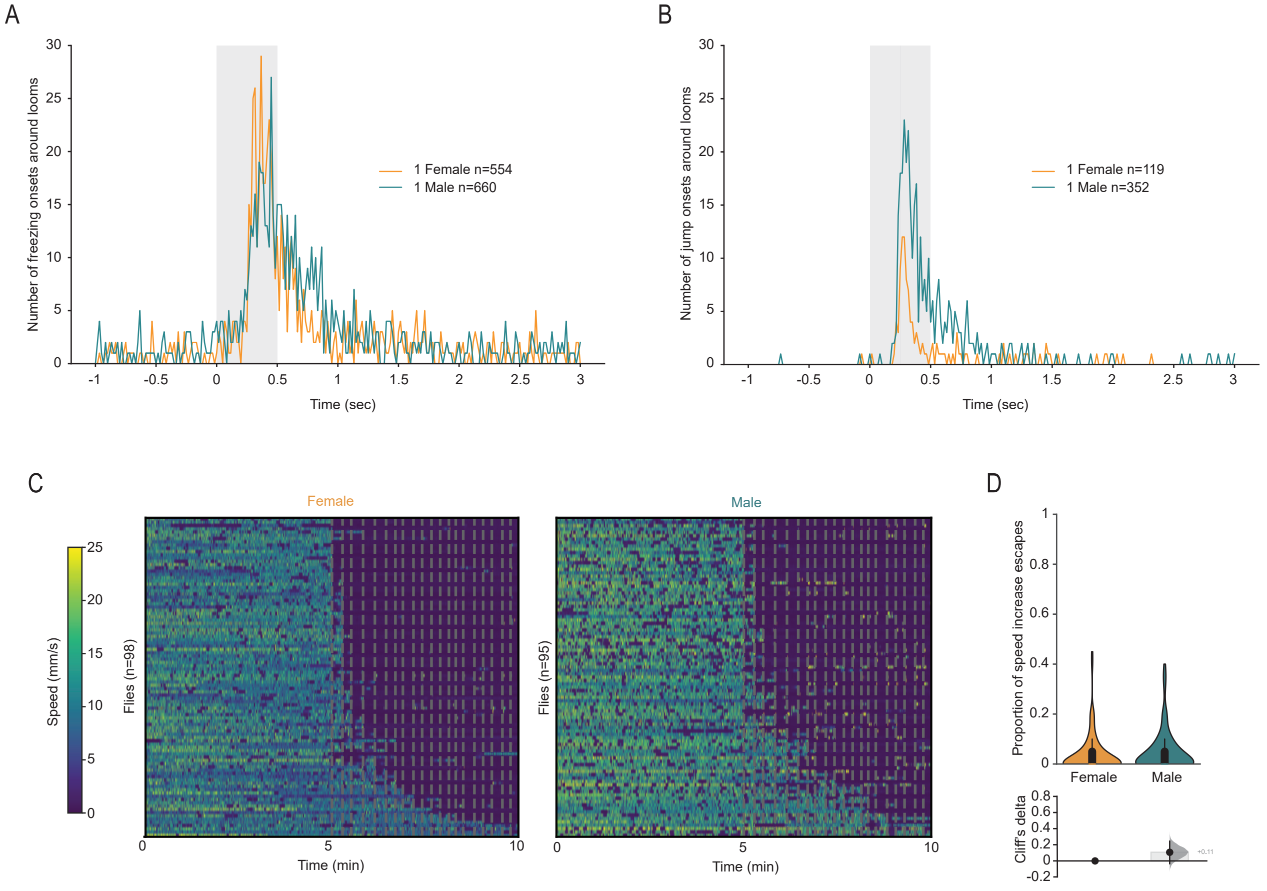


**Supplementary Figure 1. Deployment timing and frequency of individual defensive responses, and speed profiles by sex. (A, B)** Number of freezing **(A)** and jump **(B)** onset events per frame, relative to loom onset (grey shading indicates loom presentation). **(C)** Speed (colour scale on the left) for individual females (left, n=98) and male flies (right, n=95). Each row corresponds to one fly, rank-ordered by maximum time freezing during stimulation. **(D)** Proportion of speed increase escapes in response to looming stimuli. Top: Violin plots representing the probability density distribution of individual fly data bound to the range of possible values, with boxplot elements (central white dot, median; box limits, upper (75) and lower (25) quartiles; whiskers, 1.5 × interquartile range). Bottom: Cliff's delta effect size for the female–male comparison (black dot, observed Cliff's delta; grey curve, bootstrap sampling distribution; vertical line, 95% confidence interval).

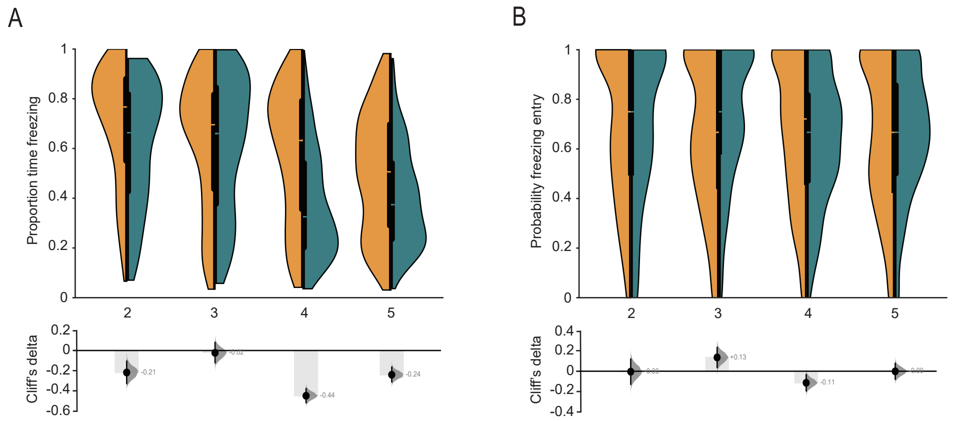


**Supplementary Figure 2. Group effect on freezing across sex. (A, B)** Proportion of time spent freezing during the stimulation **(A)** and probability of entry freezing upon loom **(B)** for females (orange) and male flies (teal). Top: Split-violin plots representing the probability density distribution of individual fly data bound to the range of possible values, with boxplot elements (central white dot, median; box limits, upper (75) and lower (25) quartiles; whiskers, 1.5 × interquartile range). Bottom: Cliff's delta effect size for the female–male comparison (black dot, observed Cliff's delta; grey curve, bootstrap sampling distribution; vertical line, 95% confidence interval).

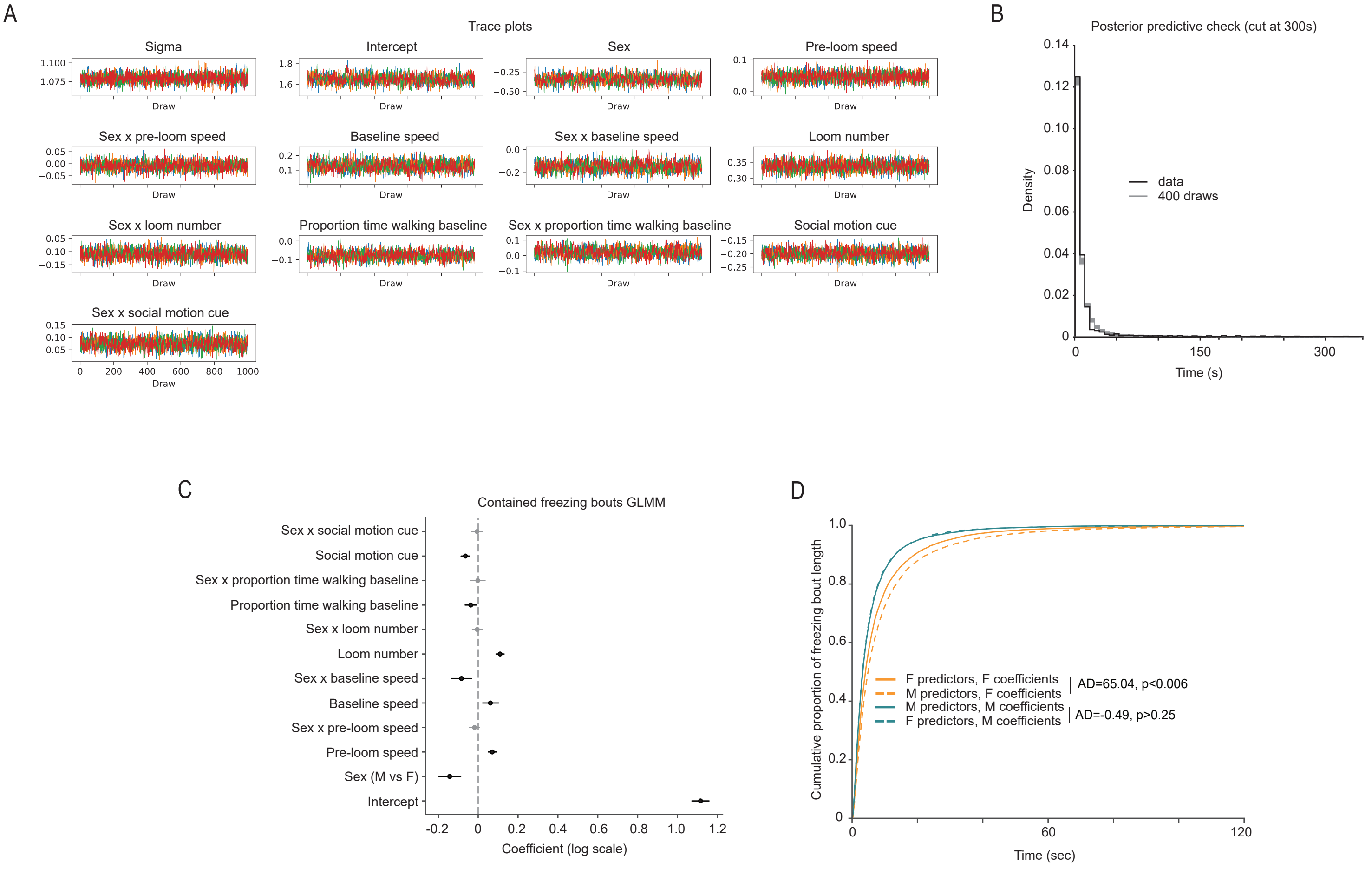


**Supplementary Figure 3. Generalised linear mixed model diagnostics and category-specific effects. (A)** Trace plots for all fixed-effect coefficients (4 chains per coefficient), showing model convergence. **(B)** Posterior predictive check (400 randomly drawn posterior samples, cut at the maximum freezing bout length, 300 s), showing model fit to the observed data. **(C)** Posterior mean coefficients (dots) and 89% credible intervals (lines; black excludes zero, grey crosses zero) for the contained freezing bout GLMMs. **(D)** Empirical cumulative distribution of simulated freeze bout durations (n = 10,000 per scenario) from the fitted lognormal model, for female and male predictor profiles combined with their own or the other sex's coefficients. Anderson-Darling (AD) test statistic and p-value are shown.

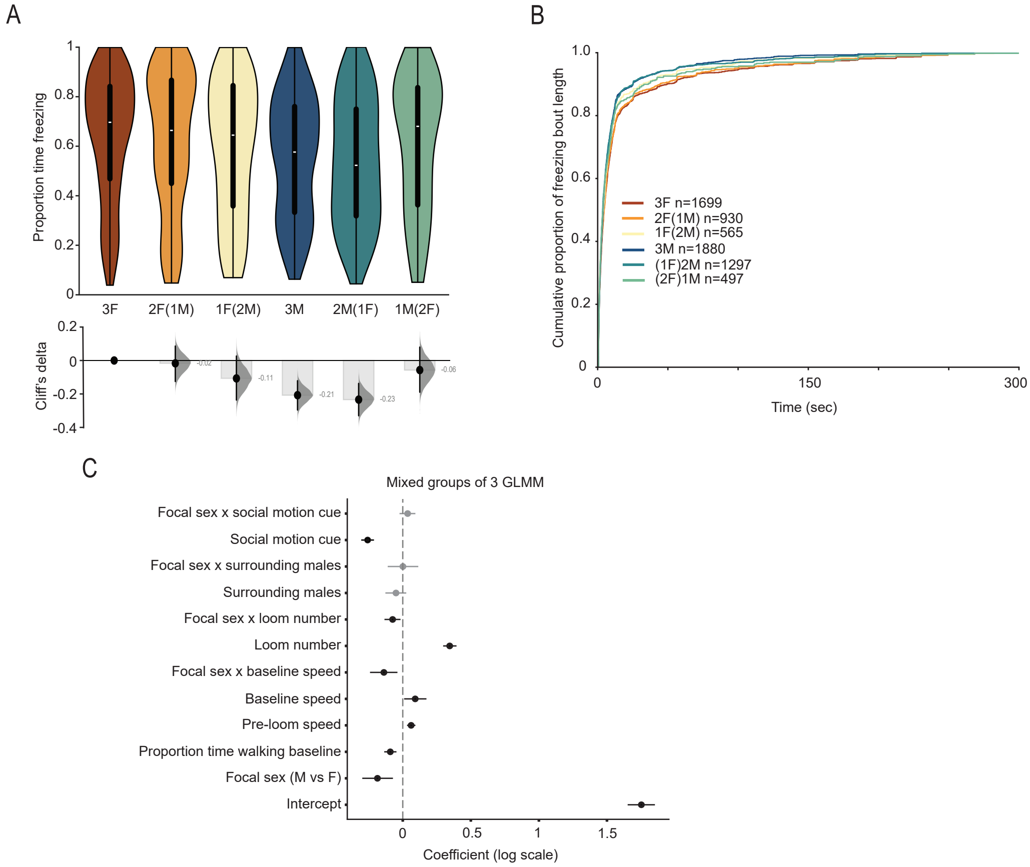


**Supplementary Figure 4. Sex differences in mixed groups of 3 flies. (A)** Proportion of time freezing during the stimulation per sex for flies tested in groups of 3 with different numbers of surrounding same sex flies. Top: Split-violin plots representing the probability density distribution of individual fly data bound to the range of possible values, with boxplot elements (central white dot, median; box limits, upper (75) and lower (25) quartiles; whiskers, 1.5 × interquartile range). Bottom: Cliff's delta effect size for the female–male comparison (black dot, observed Cliff's delta; grey curve, bootstrap sampling distribution; vertical line, 95% confidence interval). **(B)** Cumulative proportion of freezing bout length per sex for flies tested in groups of 3 with different numbers of surrounding same sex flies (see legend inset for colour code and number of freezing bouts). **(C)** Generalised linear mixed model of freeze bout duration, with a log-normal distribution, with focal sex as a categorical predictor (female as the reference level) and surrounding sex and other relevant predictors uncovered through the single sex GLMM. Posterior mean coefficients (dots) and 89% credible intervals (lines; black excludes zero; grey crosses zero).
