## Supplementary Tables for "Neuronal sex identity shapes social buffering of defensive freezing behaviour in fruit flies"

Supplementary Table 1 – Time spent freezing during looming stimulation in females: individually tested flies compared with groups of 2–5 flies.

a) Females

| Comparison | Group A (median, IQR) | Group B (median, IQR) | δ | 95% BCa CI | p | nA | nB |
|---|---|---|---|---|---|---|---|
| 1F vs 2F | 0.92, 0.80–0.96 | 0.77, 0.55–0.88 | −0.42 | [−0.54, −0.28] | <0.001 | 98 | 196 |
| 1F vs 3F | 0.92, 0.80–0.96 | 0.70, 0.44–0.82 | −0.53 | [−0.64, −0.42] | <0.001 | 98 | 252 |
| 1F vs 4F | 0.92, 0.80–0.96 | 0.63, 0.36–0.79 | −0.61 | [−0.70, −0.49] | <0.001 | 98 | 396 |
| 1F vs 5F | 0.92, 0.80–0.96 | 0.51, 0.29–0.70 | −0.74 | [−0.81, −0.63] | <0.001 | 98 | 450 |

*Values are median proportion of time spent freezing with interquartile range (IQR). δ = Cliff's delta; CI = bias-corrected and accelerated (BCa) bootstrap confidence interval; n = number of flies.*

b) Males

| Comparison | Group A (median, IQR) | Group B (median, IQR) | δ | 95% BCa CI | p | nA | nB |
|---|---|---|---|---|---|---|---|
| 1M vs 2M | 0.87, 0.69–0.94 | 0.66, 0.43–0.82 | −0.47 | [−0.59, −0.34] | <0.001 | 95 | 196 |
| 1M vs 3M | 0.87, 0.69–0.94 | 0.66, 0.38–0.85 | −0.42 | [−0.53, −0.29] | <0.001 | 95 | 255 |
| 1M vs 4M | 0.87, 0.69–0.94 | 0.33, 0.20–0.54 | −0.81 | [−0.87, −0.72] | <0.001 | 95 | 416 |
| 1M vs 5M | 0.87, 0.69–0.94 | 0.37, 0.23–0.54 | −0.79 | [−0.86, −0.70] | <0.001 | 95 | 465 |

*Values are median proportion of time spent freezing with interquartile range (IQR). δ = Cliff's delta; CI = bias-corrected and accelerated (BCa) bootstrap confidence interval; n = number of flies.*

Supplementary Table 2 – Time spent freezing during looming stimulation by group size and sex.

| Group size | PropFF, IQR | PropFM, IQR | δ | 95% BCa CI | p | nF | nM |
|---|---|---|---|---|---|---|---|
| 2 | 0.77, 0.55–0.88 | 0.66, 0.43–0.82 | −0.21 | [−0.32, −0.10] | <0.001 | 196 | 196 |

| Group size | PropFF, IQR | PropFM, IQR | δ | 95% BCa CI | p | nF | nM |
|---|---|---|---|---|---|---|---|
| 3 | 0.70, 0.44–0.82 | 0.66, 0.38–0.85 | −0.023 | [−0.12, 0.08] | 0.649 | 252 | 255 |
| 4 | 0.63, 0.36–0.79 | 0.33, 0.20–0.54 | −0.44 | [−0.51, −0.37] | <0.001 | 396 | 416 |
| 5 | 0.51, 0.29–0.70 | 0.37, 0.23–0.54 | −0.24 | [−0.31, −0.16] | <0.001 | 450 | 465 |

*Values are median proportion of time spent freezing with interquartile range (IQR). δ = Cliff's delta; CI = bias-corrected and accelerated (BCa) bootstrap confidence interval; n = number of flies.*

Supplementary Table 3 – Probability of freezing entry by group size and sex.

| Group size | PFentryF, IQR | PFentryM, IQR | δ | 95% BCa CI | p | nF | nM |
|---|---|---|---|---|---|---|---|
| 2 | 0.75, 0.50–1.00 | 0.75, 0.50–1.00 | −0.005 | [−0.13, 0.12] | 0.95 | 156 | 158 |
| 3 | 0.67, 0.44–1.00 | 0.75, 0.58–1.00 | 0.13 | [0.034, 0.23] | 0.01 | 235 | 238 |
| 4 | 0.72, 0.46–1.00 | 0.67, 0.47–0.82 | −0.11 | [−0.19, −0.03] | 0.006 | 386 | 413 |
| 5 | 0.67, 0.43–1.00 | 0.67, 0.50–0.86 | −0.001 | [−0.08, 0.08] | 0.98 | 406 | 423 |

*Values are median probability of freezing entry (PFentry) with interquartile range (IQR). δ = Cliff's delta; CI = bias-corrected and accelerated (BCa) bootstrap confidence interval; n = number of flies.*

Supplementary Table 4 – Freezing bout duration by group and sex.

| Group size | FboutF (s), IQR | FboutM (s), IQR | AD | p | nF | nM |
|---|---|---|---|---|---|---|
| 2 | 4.43, 1.40–10.88 | 3.80, 1.53–9.23 | 3.35 | 0.014 | 940 | 1237 |
| 3 | 4.47, 1.45–10.83 | 4.07, 1.45–9.55 | 2.23 | 0.039 | 1299 | 1649 |
| 4 | 4.40, 1.52–9.42 | 2.77, 1.15–5.94 | 112.67 | <0.001 | 2460 | 3643 |
| 5 | 3.35, 1.22–7.38 | 2.83, 1.22–5.70 | 37.92 | <0.001 | 3323 | 4375 |

*Values are median freeze bout duration (Fbout, s) with interquartile range (IQR). AD = Anderson-Darling test statistic; n = number of bouts.*

Supplementary Table 5 – Number of jumps per fly by group size and sex, with Cliff's delta comparison between sexes.

| Group size | NJumpF, IQR | NJumpM, IQR | δ | 95% BCa CI | p | nF | nM |
|---|---|---|---|---|---|---|---|
| 2 | 1, 0–2 | 2, 0–4 | 0.200 | [0.09, 0.31] | <0.001 | 196 | 196 |
| 3 | 0, 0–1 | 2, 0–5 | 0.418 | [0.33, 0.50] | <0.001 | 252 | 255 |
| 4 | 0, 0–1 | 1, 0–4 | 0.303 | [0.23, 0.37] | <0.001 | 396 | 416 |
| 5 | 0, 0–1 | 1, 0–3 | 0.352 | [0.29, 0.41] | <0.001 | 450 | 465 |

*Values are median number of jumps (NJump) with interquartile range (IQR). δ = Cliff's delta; CI = bias-corrected and accelerated (BCa) bootstrap confidence interval; n = number of flies.*

Supplementary Table 6 – Speed increase magnitude relative to pre-loom behaviour, by group size and sex.

| Group size | SpeedIncF (mm/s), IQR | SpeedIncM (mm/s), IQR | δ | 95% BCa CI | p | nF | nM |
|---|---|---|---|---|---|---|---|
| 2 | 2.51, 1.46–4.01 | 3.98, 2.25–6.49 | 0.34 | [0.20, 0.47] | <0.001 | 102 | 126 |
| 3 | 2.55, 1.62–3.65 | 3.93, 2.45–5.39 | 0.32 | [0.19, 0.44] | <0.001 | 154 | 153 |
| 4 | 2.35, 1.42–3.48 | 3.65, 2.48–5.38 | 0.38 | [0.29, 0.46] | <0.001 | 252 | 353 |
| 5 | 2.45, 1.63–3.58 | 3.76, 2.26–5.79 | 0.33 | [0.25, 0.40] | <0.001 | 329 | 378 |

*Values are median speed increase (SpeedInc, mm/s) with interquartile range (IQR). δ = Cliff's delta; CI = BCa bootstrap confidence interval; n = number of flies.*

Supplementary Table 7 – Proportion of speed increase occurrence, by group size and sex.

| Group size | PropSpeedIncF, IQR | PropSpeedIncM, IQR | δ | 95% BCa CI | p | nF | nM |
|---|---|---|---|---|---|---|---|
| 2 | 0.05, 0–0.10 | 0.05, 0–0.15 | 0.17 | [0.058, 0.27] | 0.004 | 196 | 196 |
| 3 | 0.05, 0–0.15 | 0.05, 0–0.15 | −0.023 | [−0.12, 0.072] | 0.64 | 252 | 255 |
| 4 | 0.05, 0–0.15 | 0.10, 0.05–0.20 | 0.31 | [0.24, 0.38] | <0.001 | 396 | 416 |

| Group size | PropSpeedIncF, IQR | PropSpeedIncM, IQR | δ | 95% BCa CI | p | nF | nM |
|---|---|---|---|---|---|---|---|
| 5 | 0.10, 0–0.19 | 0.10, 0.05–0.20 | 0.092 | [0.019, 0.17] | 0.015 | 450 | 465 |

*Values are median probability of speed increase occurrence (PropSpeedInc) with interquartile range (IQR). δ = Cliff's delta; CI = BCa bootstrap confidence interval; n = number of flies.*

Supplementary Table 8 – Comparison of candidate error distributions for the main log-normal GLMM (groups, pooled sizes 2–5), via leave-one-out cross-validation (LOO-CV).

| Model | Rank | ΔELPD | SE (Δ) | Pareto $\hat{k}$ > 0.7 (n) | Weight |
|---|---|---|---|---|---|
| Log-normal | 0 | 0 | 0 | 1 | 1.00 |
| Weibull | 1 | −1500 | 45 | 437 | 0.00 |
| Gamma | 2 | −1500 | 50 | 493 | 0.00 |

*ΔELPD (expected log predictive density) and its standard error are reported relative to the best-supported model (log-normal, rank 0). Pareto $\hat{k}$ > 0.7 indicates the number of observations for which the LOO importance-sampling approximation is unreliable. Weight reflects the relative predictive support (stacking weight) assigned to each model.*

Supplementary Table 9 – Posterior estimates and convergence diagnostics for the main log-normal GLMM (groups, pooled sizes 2–5).

| Term | Mean | 89% CI | R-hat | ESS bulk | ESS tail |
|---|---|---|---|---|---|
| Intercept | 1.65 | [1.58, 1.72] | 1.002 | 1169 | 1464 |
| Sex (M vs F) | −0.34 | [−0.43, −0.25] | 1.000 | 1195 | 1880 |
| Pre-loom speed | 0.04 | [0.02, 0.06] | 1.001 | 3378 | 2815 |
| Sex × pre-loom speed | −0.009 | [−0.038, 0.020] | 1.001 | 3364 | 3068 |
| Baseline speed | 0.13 | [0.076, 0.18] | 1.003 | 1377 | 2104 |
| Sex × baseline speed | −0.15 | [−0.21, −0.083] | 1.004 | 1342 | 2035 |
| Loom number | 0.34 | [0.31, 0.36] | 1.001 | 2806 | 2799 |
| Sex × loom number | −0.11 | [−0.14, −0.080] | 1.002 | 3140 | 2847 |
| Proportion time walking (baseline) | −0.077 | [−0.11, −0.040] | 1.003 | 1522 | 1997 |
| Sex × proportion time walking | 0.020 | [−0.027, 0.069] | 1.004 | 1554 | 1869 |
| Social motion cue | −0.20 | [−0.23, −0.17] | 1.001 | 2609 | 2413 |

| Term | Mean | 89% CI | R-hat | ESS bulk | ESS tail |
|---|---|---|---|---|---|
| Sex × social motion cue | 0.075 | [0.041, 0.11] | 1.001 | 2577 | 2363 |
| sigma (dispersion) | 1.079 | [1.070, 1.089] | 1.001 | 3387 | 2658 |

*Mean and 89% credible interval (CI) of the posterior distribution for each fixed effect, on the log scale. R-hat (Gelman–Rubin potential scale reduction factor) values indicate convergence across chains. ESS bulk and ESS tail report the effective sample size.*

Supplementary Table 10 – Breakdown of contained versus extended freeze bouts by group size and sex.

| Group size | Sex | Contained (n) | Extended (n) | Total (n) |
|---|---|---|---|---|
| 2 | F | 770 | 170 | 940 |
| 2 | M | 1059 | 178 | 1237 |
| 3 | F | 1075 | 224 | 1299 |
| 3 | M | 1402 | 247 | 1649 |
| 4 | F | 2109 | 351 | 2460 |
| 4 | M | 3416 | 227 | 3643 |
| 5 | F | 3007 | 316 | 3323 |
| 5 | M | 4149 | 226 | 4375 |

*Contained bouts resolved before the onset of the next loom; extended bouts persisted into a subsequent inter-loom interval. n = number of bouts.*

Supplementary Table 11 – Pairwise comparison of simulated freeze bout durations in the counterfactual sex-swap analysis of the main log-normal GLMM.

| Comparison | Median A (s) | Median B (s) | Δ median (A − B), s [95% CI] | AD | p (adj.) | nA | nB |
|---|---|---|---|---|---|---|---|
| F profile + F coefs vs M profile + M coefs | 4.64 | 3.78 | 0.86 [0.70, 1.01] | 111.05 | <0.006 | 10000 | 10000 |
| F profile + F coefs vs M profile + F coefs | 4.64 | 5.52 | −0.88 [−1.05, −0.71] | 65.04 | <0.006 | 10000 | 10000 |
| F profile + F coefs vs F profile + M coefs | 4.64 | 3.73 | 0.90 [0.76, 1.05] | 126.30 | <0.006 | 10000 | 10000 |
| M profile + M coefs vs M profile + F coefs | 3.78 | 5.52 | −1.74 [−1.89, −1.58] | 347.01 | <0.006 | 10000 | 10000 |
| M profile + M coefs vs F profile + M coefs | 3.78 | 3.73 | 0.05 [−0.08, 0.17] | −0.49 | >0.25 | 10000 | 10000 |
| M profile + F coefs vs F profile + M coefs | 5.52 | 3.73 | 1.79 [1.64, 1.94] | 374.24 | <0.006 | 10000 | 10000 |

*Freeze bout durations were simulated from the main log-normal GLMM for each scenario. 'Profile' denotes the locomotion values (baseline and pre-loom speeds and social motion cues) of the sex indicated and 'coefs' the coefficients of that sex (weight of each predictor and its interactions). Values are median simulated bout duration (s). Δ = difference in medians (group A − group B) with 95% confidence interval; AD = Anderson-Darling k-sample test statistic; n = number of simulated bouts. p-values are Bonferroni-corrected (k = 6); where the uncorrected AD p-value reached the tabulated floor (p < 0.001), the corrected value is reported as an upper bound, and where it reached the cap (p ≥ 0.25) the corrected value is 1.000.*

Supplementary Table 12 – Time spent freezing in mixed-sex groups of three.

| Comparison | PropFreeze, group A, IQR | PropFreeze, group B, IQR | δ | 95% BCa CI | p (adj.) | nA | nB |
|---|---|---|---|---|---|---|---|
| 3F vs 2F1M (F) | 0.70, 0.47–0.84 | 0.66, 0.45–0.87 | −0.017 | [−0.13, 0.09] | 1.000 | 333 | 194 |
| 3F vs 1F2M (F) | 0.70 0.47–0.84 | 0.64, 0.36–0.85 | −0.11 | [−0.24, 0.027] | 0.526 | 333 | 99 |

| Comparison | PropFreeze, group A, IQR | PropFreeze, group B, IQR | δ | 95% BCa CI | p (adj.) | nA | nB |
|---|---|---|---|---|---|---|---|
| 3F vs 3M | 0.70, 0.47–0.84 | 0.58, 0.33–0.76 | −0.21 | [−0.30, −0.12] | <0.001 | 333 | 291 |
| 3F vs 1F2M (M) | 0.70, 0.47–0.84 | 0.52, 0.32–0.75 | −0.23 | [−0.33, −0.14] | <0.001 | 333 | 198 |
| 3F vs 2F1M (M) | 0.70, 0.47–0.84 | 0.68, 0.36–0.84 | −0.057 | [−0.19, 0.08] | 1.000 | 333 | 97 |

*Values are median proportion of the time spent freezing in the stimulation (PropFreeze) with interquartile range (IQR). δ = Cliff's delta for group B relative to group A; CI = BCa bootstrap confidence interval; n = number of flies.*

Supplementary Table 13 – Freeze bout duration by sex and across sex in mixed-sex groups of 3.

| Comparison | Group A (median, IQR) | Group B (median, IQR) | AD | p (adj.) | nA | nB |
|---|---|---|---|---|---|---|
| 3F vs 3M | 4.53, 1.52–11.47 | 3.93, 1.43–9.23 | 11.47 | <0.009 | 1693 | 1871 |
| 2F1M(F) vs 1F2M(M) | 4.63, 1.38–11.78 | 3.55, 1.33–9.02 | 6.74 | <0.009 | 927 | 1294 |
| 1F2M(F) vs 2F1M(M) | 4.13, 1.18–9.96 | 3.82, 1.37–10.00 | −0.57 | 1.000 | 559 | 497 |
| 3F vs 2F1M(F) | 4.53, 1.52–11.47 | 4.63, 1.38–11.78 | 0.15 | 1.000 | 1693 | 927 |
| 3F vs 1F2M(F) | 4.53, 1.52–11.47 | 4.13, 1.18–9.96 | 2.87 | 0.198 | 1693 | 559 |
| 2F1M(F) vs 1F2M(F) | 4.63, 1.38–11.78 | 4.13, 1.18–9.96 | 0.88 | 1.000 | 927 | 559 |
| 3M vs 1F2M(M) | 3.93, 1.43–9.23 | 3.55, 1.33–9.02 | −0.02 | 1.000 | 1871 | 1294 |
| 3M vs 2F1M(M) | 3.93, 1.43–9.23 | 3.82, 1.37–10.00 | 0.30 | 1.000 | 1871 | 497 |
| 1F2M(M) vs 2F1M(M) | 3.55, 1.33–9.02 | 3.82, 1.37–10.00 | −0.25 | 1.000 | 1294 | 497 |

*Values are median freeze bout duration (s) with interquartile range (IQR). AD = Anderson-Darling test statistic; n = number of bouts. p-values are Bonferroni-corrected across the nine comparisons shown (k = 9). Where the uncorrected Anderson-Darling p-value reached the tabulated floor (p < 0.001) or ceiling (p = 0.25), the corrected value is reported as an upper or lower bound accordingly.*

Supplementary Table 14 – Posterior estimates and convergence diagnostics for the log-normal GLMM (mixed-sex groups of 3).

| Term | Mean | 89% CI | R-hat | ESS bulk | ESS tail |
|---|---|---|---|---|---|
| Intercept | 1.75 | [1.66, 1.85] | 1.004 | 1090 | 1501 |

| Term | Mean | 89% CI | R-hat | ESS bulk | ESS tail |
|---|---|---|---|---|---|
| Focal sex (M vs F) | −0.19 | [−0.29, −0.077] | 1.002 | 1488 | 2252 |
| Surrounding males | −0.050 | [−0.12, 0.021] | 1.001 | 965 | 1311 |
| Pre-loom speed | 0.062 | [0.036, 0.088] | 0.999 | 4601 | 3411 |
| Focal sex × baseline speed | −0.14 | [−0.24, −0.044] | 1.002 | 1605 | 2286 |
| Baseline speed | 0.091 | [0.015, 0.17] | 1.002 | 1387 | 1814 |
| Loom number | 0.34 | [0.30, 0.39] | 1.003 | 2116 | 2133 |
| Focal sex × loom number | −0.075 | [−0.13, −0.022] | 1.002 | 2546 | 2773 |
| Proportion time walking (baseline) | −0.092 | [−0.13, −0.051] | 1.000 | 2139 | 2727 |
| Social motion cue | −0.26 | [−0.30, −0.22] | 1.002 | 2060 | 2557 |
| Focal sex × social motion cue | 0.036 | [−0.019, 0.088] | 1.003 | 2223 | 2682 |
| Focal sex × surrounding males | 0.001 | [−0.11, 0.11] | 1.002 | 956 | 1888 |
| sigma (dispersion) | 1.24 | [1.22, 1.26] | 1.000 | 3428 | 3193 |

*Mean and 89% credible interval (CI) of the posterior distribution for each fixed effect, on the log scale. R-hat (Gelman–Rubin potential scale reduction factor) values indicate convergence across chains. ESS bulk and ESS tail report the effective sample size.*

Supplementary Table 15 – Time spent freezing in the nervous system sex-reversal experiment: individually tested flies compared with groups of 5.

| Comparison | PropFreeze, group A, IQR | PropFreeze, group B, IQR | δ | 95% BCa CI | p (adj.) | nA | nB |
|---|---|---|---|---|---|---|---|
| Elav-GAL4/+: 1 vs 5 flies | 0.73, 0.37–0.91 | 0.20, 0.14–0.27 | −0.68 | [−0.79, −0.54] | <0.001 | 102 | 500 |
| UAS-tra/+: 1 vs 5 flies | 0.75, 0.34–0.94 | 0.20, 0.14–0.27 | −0.63 | [−0.74, −0.49] | <0.001 | 102 | 510 |
| Elav-GAL4>UAS-Tra (feminised): 1 vs 5 flies | 0.72, 0.44–0.89 | 0.37, 0.22–0.62 | −0.42 | [−0.54, −0.28] | <0.001 | 102 | 510 |

*Group A = individually tested flies and group B = groups of 5 flies (males). Values are median proportion of the time spent freezing in the stimulation (PropFreeze) with interquartile range (IQR). δ = Cliff's delta for group B relative to group A; CI = BCa bootstrap confidence interval; p-values are Bonferroni-corrected (k = 3); n = number of flies.*

Supplementary Table 16 – Freeze bout duration in the nervous system sex-reversal experiment: group-tested flies (groups of 5).

| Comparison | Fbout, group A (s), IQR | Fbout, group B (s), IQR | AD | p (adj.) | nA | nB |
|---|---|---|---|---|---|---|
| Elav-GAL4/+ vs UAS-Tra/+ | 1.55, 0.85–3.08 | 1.50, 0.87–2.93 | 0.94 | 0.402 | 3657 | 3588 |
| Elav-GAL4 vs Elav-GAL4>UAS-Tra (feminised) | 1.55, 0.85–3.08 | 2.10, 0.98–4.85 | 110.49 | <0.003 | 3657 | 3115 |
| UAS-Tra/+ vs Elav-GAL4>UAS-Tra (feminised) | 1.50, 0.87–2.93 | 2.10, 0.98–4.85 | 116.68 | <0.003 | 3588 | 3115 |

*Values are median freeze bout duration (Fbout, s) with interquartile range (IQR). AD = Anderson-Darling test statistic; n = number of bouts. p-values are Bonferroni-corrected (k = 3); where the uncorrected Anderson-Darling p-value reached the tabulated floor (p < 0.001), the corrected value is reported as an upper bound.*

Supplementary Table 17 – Time spent freezing in the LC11 silencing experiment (Empty-GAL4>TNT vs LC11-GAL4>TNT): individually tested flies compared with groups of 5, by genotype and sex.

| Comparison | PropFreeze, group A, IQR | PropFreeze, group B, IQR | δ | 95% BCa CI | p | nA | nB |
|---|---|---|---|---|---|---|---|
| Empty-GAL4>TNT (F): 1 vs 5 flies | 0.79, 0.41–0.95 | 0.19, 0.08–0.44 | −0.56 | [−0.67, −0.43] | <0.001 | 103 | 330 |
| Empty-GAL4>TNT (M): 1 vs 5 flies | 0.27, 0.08–0.71 | 0.12, 0.07–0.22 | −0.31 | [−0.45, −0.16] | <0.001 | 103 | 390 |
| LC11-GAL4>TNT (F): 1 vs 5 flies | 0.76, 0.39–0.93 | 0.72, 0.44–0.90 | −0.032 | [−0.16, 0.10] | 0.620 | 104 | 375 |
| LC11-GAL4>TNT (M): 1 vs 5 flies | 0.70, 0.20–0.89 | 0.64, 0.39–0.84 | −0.024 | [−0.16, 0.12] | 0.700 | 104 | 390 |

*Group A = individually tested flies and group B = groups of 5 flies. Values are median proportion of the time spent freezing in the stimulation (PropFreeze) with interquartile range (IQR). δ = Cliff's delta for group B relative to group A; CI = BCa bootstrap confidence interval; n = number of flies.*

Supplementary Table 18 – Freeze bout duration in the LC11 silencing experiment (Empty-GAL4>TNT vs LC11-GAL4>TNT): group-tested flies (groups of 5).

| Comparison | Fbout, group A (s), IQR | Fbout, group B (s), IQR | AD | p | nA | nB |
|---|---|---|---|---|---|---|
| Empty-GAL4>TNT vs LC11-GAL4>TNT (F) | 2.29, 0.97–5.85 | 3.10, 1.20–9.80 | 32.19 | <0.001 | 1420 | 1213 |
| Empty-GAL4>TNT vs LC11-GAL4>TNT (M) | 1.90, 0.87–4.49 | 2.73, 1.07–7.76 | 57.68 | <0.001 | 1480 | 1304 |

*Values are median freeze bout duration (Fbout, s) with interquartile range (IQR). AD = Anderson-Darling test statistic; n = number of bouts.*

Supplementary Table 19 – Freeze bout duration in the LC11 silencing experiment (Empty-GAL4>TNT vs LC11-GAL4>TNT): comparison between females and males in group-tested flies (groups of 5).

| Comparison | Fbout F (s), IQR | Fbout M (s), IQR | AD | p | nF | nM |
|---|---|---|---|---|---|---|
| Empty-GAL4>TNT: females vs males | 2.29, 0.97–5.85 | 1.90, 0.87–4.49 | 16.75 | <0.001 | 1420 | 1480 |
| LC11-GAL4>TNT: females vs males | 3.10, 1.20–9.80 | 2.73, 1.07–7.76 | 3.46 | 0.013 | 1213 | 1304 |

*Values are median freeze bout duration (Fbout, s) with interquartile range (IQR). AD = Anderson-Darling test statistic; n = number of bouts.*
